# High-resolution radioluminescence microscopy of FDG uptake in an engineered 3D tumor-stoma model

**DOI:** 10.1101/2020.12.07.412700

**Authors:** Syamantak Khan, Sungwoo Kim, Yunzhi Peter Yang, Guillem Pratx

## Abstract

**PURPOSE:** The increased glucose metabolism of cancer cells is the basis for ^18^F-fluorodeoxyglucose positron emission tomography (FDG-PET). However, due to its coarse image resolution, PET is unable to resolve the metabolic role of cancer-associated stroma, which often influences the metabolic reprogramming of a tumor. This study investigates the feasibility of imaging engineered 3D tumor models with high resolution using FDG.

**METHOD:** Multicellular tumor spheroids (A549 lung adenocarcinoma) were co-cultured with GFP-expressing human umbilical vein endothelial cells (HUVECs) within an artificial extracellular matrix to mimic a tumor and its surrounding stroma. The tumor model was constructed as a thin 3D layer over a transparent CdWO4 scintillator plate to allow high-resolution imaging of the cultured cells. The radioluminescence signal was collected by a highly sensitive microscope and camera. Fluorescence microscopy was performed in the same instrument to localize endothelial and tumor cells.

**RESULTS:** Simultaneous brightfield and fluorescence imaging, co-localized with radioluminescence signal, provided high-resolution information on FDG accumulation in the engineered tissue. The microvascular stromal compartment took up a large fraction of FDG, comparable to the tumor spheroids. In vitro gamma counting also revealed that both A549 and HUVEC cells were highly glycolytic and characterized by rapid FDG-uptake kinetics.

**CONCLUSION:** Our study demonstrates the feasibility of imaging FDG distribution in tumor and stromal components separately with high spatial resolution in engineered *in vitro* tumor models. Our imaging method is safe, simple, rapid, and can be easily used for other *in vitro* cancer models, surgical tissue slices, and tumor biopsies to interrogate PET radiotracer uptake at high resolution.

## Introduction

Cancer cells in most tumors are glycolytic, characterized by a higher rate of glucose uptake compared to the normal surrounding tissue, even when sufficient oxygen is present for oxidative glucose metabolism [1]. This phenomenon of aerobic glycolysis has led to the development and widespread use of ^18^F-fluorodeoxyglucose positron emission tomography (FDG-PET) for tumor imaging, which has been shown to significantly improve the diagnosis, staging, and subsequent treatment of cancers. However, the spatial resolution of positron emission tomography (PET), which is in the range of 4-8 mm [2], creates a bottleneck for its application to complex and heterogeneous targets.

This range of imaging resolution is insufficient for resolving the structural and functional components of a tumor microenvironment, which plays a pivotal role in tumorigenesis, tumor-homeostasis, and cancer metastasis. The stromal compartment of a tumor consists of connective tissue cells that support the function of the cancerous cells. Generally, any cancer-associated stroma is enriched with normal and cancer-associated fibroblasts, blood-vessel-forming endothelial cells, and immune cells including T-cells and macrophages. How the metabolism of the stroma is functionally reprogrammed to support the cancer cells is not entirely well understood. However, it is likely that the stroma contributes significantly to the tumor uptake value measured by FDG-PET imaging. Glycolysis is significantly upregulated in proliferating fibroblasts and endothelial cells [3–5], thus one would expect these cells to take up FDG and contribute to the total tumor signal measured by PET imaging. Similarly, anti-inflammatory tumor-associated macrophages, which promote tumor-associated angiogenesis and immunosuppression by altering metabolism in breast cancer cells, [6] may also contribute to the FDG signal.

In this context, we previously demonstrated the feasibility of imaging PET tracers *in vitro* in cell cultures with single-cell resolution, using an approach known as radioluminescence microscopy (RLM) [7–10]. RLM functions by converting the emitted positrons from a radioactive sample into photons (scintillation), which are captured by a highly sensitive EMCCD camera. The technique also requires a special-purpose widefield microscopy set-up for low-light imaging and a thin scintillator plate for scintillation detection. As the average range of the emitted positrons in PET is often substantial (~1mm), high-resolution images can only be imaged by reducing the physical distance between the scintillator plate and the radioactive sample. This approach was successfully employed to image the uptake of various radiotracers by individual cells cultured directly on 2D on scintillator plates [7, 9].

Here, we demonstrate that RLM can be successfully extended for the purpose of imaging thicker tissue constructs, such as engineered tumor spheroids and stroma co-cultures. In previous studies, RLM was used to investigate PET tracers at the cellular level in 2D cell cultures, but these cultures have limited applicability since they do not accurately model the complex 3D environment in which tumor cells grow. In this study, we test the feasibility of imaging FDG uptake within a thick tissue-engineered tumor microenvironment, with high-resolution. To model a glycolytic tumor and its associated stroma, we generated A549 lung adenocarcinoma spheroids and co-cultured them with a pre-formed endothelial cell network within an artificial extracellular matrix. Using high-resolution RLM, we observed detectable FDG uptake in both the tumor and stromal compartment of this tissue-engineered model. This result demonstrates that multicellular tumor models may be suitable for *in vitro* investigation of PET tracers using RLM. Unlike other *in vitro* metabolic imaging approaches, RLM allows researchers to use the same biomarker (e.g. FDG) as PET/CT studies for rapid translation of knowledge from the pre-clinical to the clinical arena. The approach could be particularly valuable for PET tracers that target tumor-associated biomarkers expressed not by tumor cells but by stromal or immune cells.

## Materials and Methods

### Cell Culture

A549 lung adenocarcinoma cells were obtained from ATCC and cultured in standard DMEM medium with 10% fetal bovine serum, 4 mM L-glutamine, 1 mM sodium pyruvate, 100 IU penicillin, and 100 μg/ml streptomycin. To generate 3D spheroids, 2D-cultured A549 cells (50-75% confluent) were harvested by trypsinization, counted by a hemocytometer, and seeded into ultra-low-attachment round-bottom 96-well plates (Spheroid Microplates, Corning) at a density of 5,000 or 10,000 cells per well.

Immortalized human umbilical vein endothelial cells (HUVECs) expressing green fluorescent protein (GFP) were provided from the laboratory of the late Dr. J. Folkman (Children’s Hospital, Boston). HUVECs were cultured in EBM-2 (Lonza, Walkersville, MD) containing supplements from the EGM-2 kit, 10 % fetal bovine serum (FBS), and 1 % penicillin-streptomycin glutamine (PSG). The cells were cultured in an incubator supplied with 5 % CO2 at 37°C. The culture medium was changed every 3 days.

### Preparation of Gelatin Methacrylate (GelMA)

GelMA was prepared using a previous protocol [11]. Briefly, a 10 % (w/v) gelatin solution was prepared by stirring 10 g of gelatin type-A powder in 100 ml of ultra-pure distilled water and reacted at 50 °C for an hour. Then, 7.5 % (v/v) methacrylic anhydride (MA) was added to the gelatin solution dropwise, and the reaction continued at 50 °C for 6 hours. The mixture was dialyzed in distilled water using dialysis tubing (molecular weight cut off: 13,000-14,000 Da) for 6 days. After dialysis, the obtained solution was filtered and freeze-dried for 2-3 days and then stored at −20 °C until use.

### Co-culture of A549 and HUVEC cells

The GelMA hydrogel solution was prepared by mixing 10 % (w/v) GelMA and 0.1 % (w/v) of lithium phenyl(2,4,6-trimethylbenzoyl) phosphinate (LAP) in sterile PBS. The solution was sterile-filtered before mixing with cells. Cancer spheroids (day-3 after seeding 10,000 cells/spheroid or 5,000 cells/spheroid) were mixed 400,000 HUVECs in 100 ul of GelMA solution. The GelMA prepolymer solution containing cells was photo-crosslinked by visible light exposure for 1.5 minutes to create a thin hydrogel coating layer. The cell containing GelMA was transferred into a 24-well plate and cultured for 4 days. Morphological cell changes were observed using a Zeiss Axiovert 200 microscope and imaged with Zeiss Axiovision software. Fluorescent images of the cells were taken on day 0 and day 3.

### FDG labeling and sample preparation

For imaging of glucose metabolism using RLM, the hydrogel-cell mixture was cast on a CdWO_4_ scintillator plate (0.5 mm thick) to form a thin layer. After 5 days, the co-culture (and mono-cultures) were incubated in glucose-free DMEM medium for 1 h before the addition of FDG (1 mCi/ml). FDG was produced at the Stanford radiochemistry facility using an on-site cyclotron. The culture was incubated with FDG for 1-2 h followed by washing with phosphate buffer saline (PBS) thrice with gentle rocking. The CdWO_4_ scintillator plates were taken out from the 6-well plates and mounted onto a standard glass coverslip (0.1 mm thick, Fisher Scientific) upside down. The specimens were then placed on the microscope stage for imaging.

### Instrumentation and multimodal imaging

The set-up was mounted on a custom-built wide-field microscope equipped with a short focal tube lens (50 mm, 4X/0.2 NA; Nikon, CFI Plan Apochromat λ), 20X/0.75 NA air objective (Nikon, CFI Plan Apochromat λ), and deep-cooled electron-multiplying charge-coupled device (EM-CCD; Hamamatsu Photonics, ImagEM C9100-13). Brightfield images were acquired with no EM gain used as a reference focus. A 469/35 nm filter set (Thorlabs, filter ref: MDF-GFP1) was used for GFP (green) imaging of HUVEC cells. The imaging field of view was 1.5 mm with an image pixel size of 3.2 μm. For RLM imaging, images were taken with a 10X or 20X objective, an exposure time of 10-300s, an EM gain of 600/1200, and 1×1 pixel binning. Although high-quality digital imaging can be achieved for 2D cell cultures by constructing an image from individual detected events [12], the high count-rate from 3D tissue constructs creates enough scintillation signal for fast and direct analog measurements of the whole sample using a single short camera exposure. Tumor cells labeled with NucRed™ Live 647 Reagent were imaged using a 685/40 nm emission filter in a commercial fluorescence microscope (EVOS imaging system).

### Quantification of FDG concentration

To estimate the relative FDG concentration between A549 spheroids (tumor) and the HUVEC network (stroma), the EMCCD radioluminescence signal was calibrated using a known amount of FDG concentration with known radioactivity. The camera signal was related linearly to the radioactivity of FDG. The spatial concentration of radioactivity in the co-culture was obtained in Bq/pixel from the calibration curve. Line and radial distribution profiles of the radioluminescence signal were analyzed using NIH ImageJ software. For kinetic measurement, cellular FDG uptake was calculated by measuring the total radioactivity in a gamma counter and dividing by the number of cells.

## Results

### Characterization of a co-culture model of tumor and stroma

The A549 spheroids were formed within 3-4 days of seeding and showed a consistent spherical morphology (Figure 1a, 1b). Using a fluorescent hypoxia sensor (image-iT red hypoxia), we found the center of these spheroids to be hypoxic (Figure 1c). The bigger spheroids (>5,000 cells/spheroid) were flatter than the smaller ones, which retained a more spherical shape. As a necrotic core usually develops within 1-2 weeks of spheroid culture, all the spheroids were used at an earlier stage, while most of the cells were still viable. The HUVECs are known to naturally form angiogenic sprouts when grown in GelMA [13]. When HUVECs were co-cultured with A549 lung cancer spheroids, elongation, branching, and tubular networks were observed at day-3, while spheroid cancer cells largely maintained their morphologies. Migration and multiple protrusions of HUVECs were also observed into the 3D spheroids (Figure 1d-1f).

**Figure 1:**
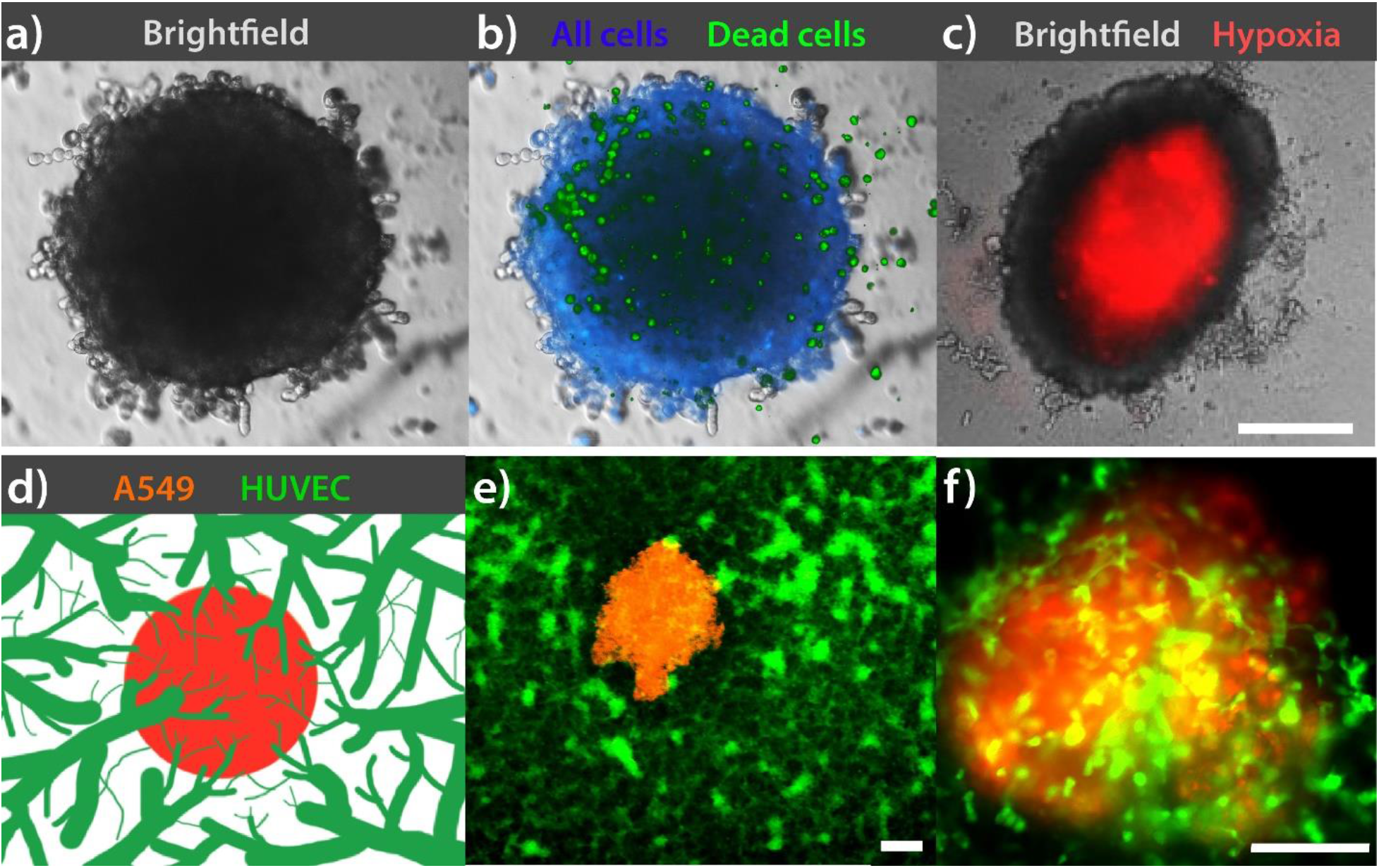
Characterization of A549 spheroids and HUVEC microvascular networks. (a) Morphology, (b) cell-viability, and (C) hypoxic core of A549 spheroid. Blue: Hoechst 33342; Green: SYTOX™ green; Red: Image-iT™ Red Hypoxia Reagent (d) Schematic and (e-f) fluorescence microscopy of the co-culture of the spheroid with an endothelial network of GFP-expressing (green) HUVECs in a hydrogel matrix. Green: Constitutive GFP expression; Orange: NucRed™ Live 647 Reagent. Scale bar: 200 μm

### Differential FDG uptake between A549 and HUVEC cells

The principle of RLM imaging and the multimodal microscopy set-up is shown in Figure 2. The scintillation light from emitted positrons traveling into the scintillator was collected using a 10X objective lens and subsequently focused on the EMCCD detector using a 0.2 NA, 50 mm imaging objective. The cells were cultured in a thin layer of a GelMA (approximately <500μm), which allowed the emitted positrons to reach the scintillator for efficient RLM imaging (Figure 2, right panel). HUVECs formed microvascular networks in GelMA and accumulated sufficient FDG to be imaged with a relatively short (~60 s) exposure (Figure 3a, b). The distribution of FDG in the microvascular network co-localized with the green fluorescence signal from GFP expression in the cells.

**Figure 2:**
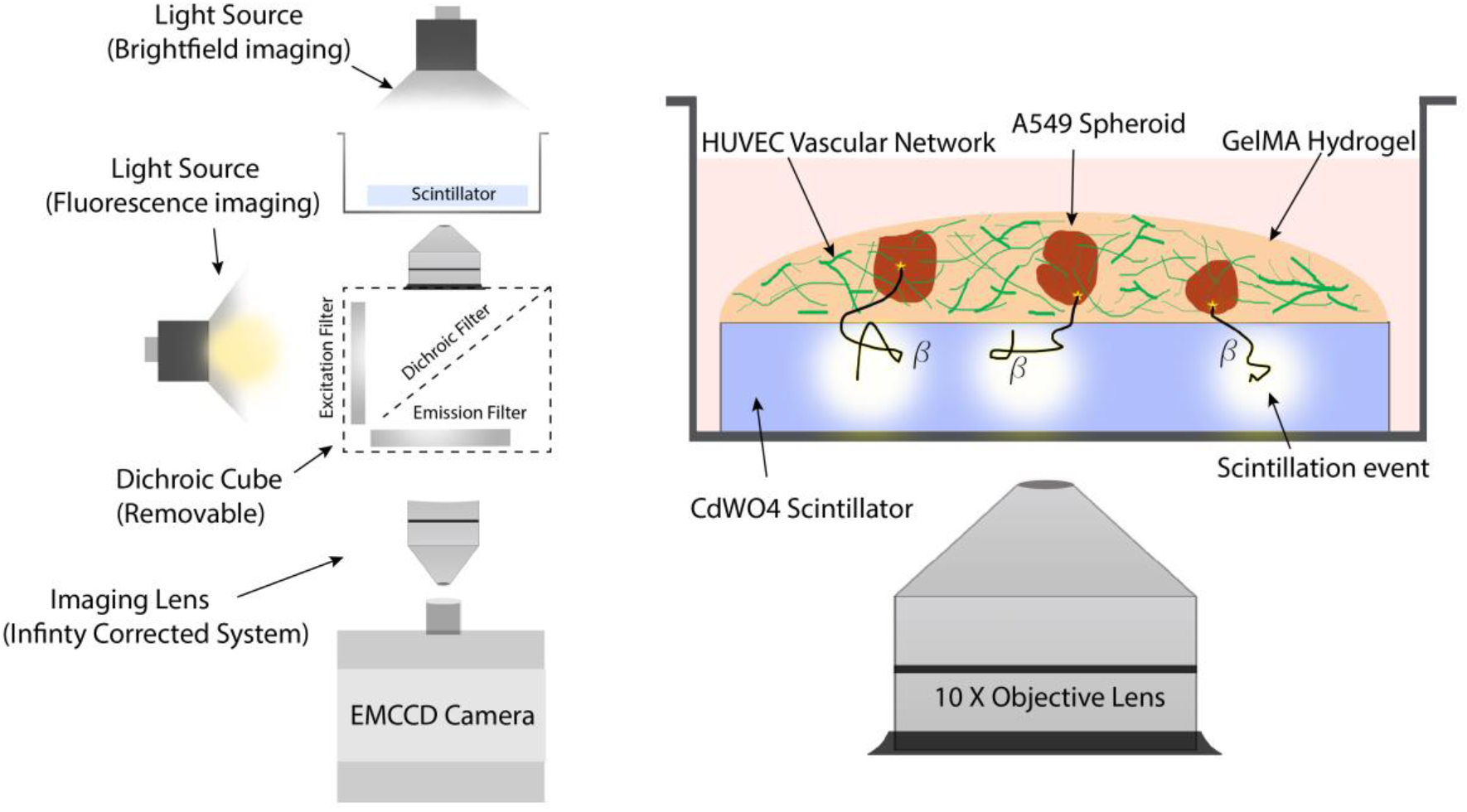
Schematic diagram of the imaging methodology used in this study. The assembly of the RLM imaging system is shown on the left. The magnified view in the right panel shows the co-culture set-up and illustrates the working principle of radioluminescence imaging.

**Figure 3:**
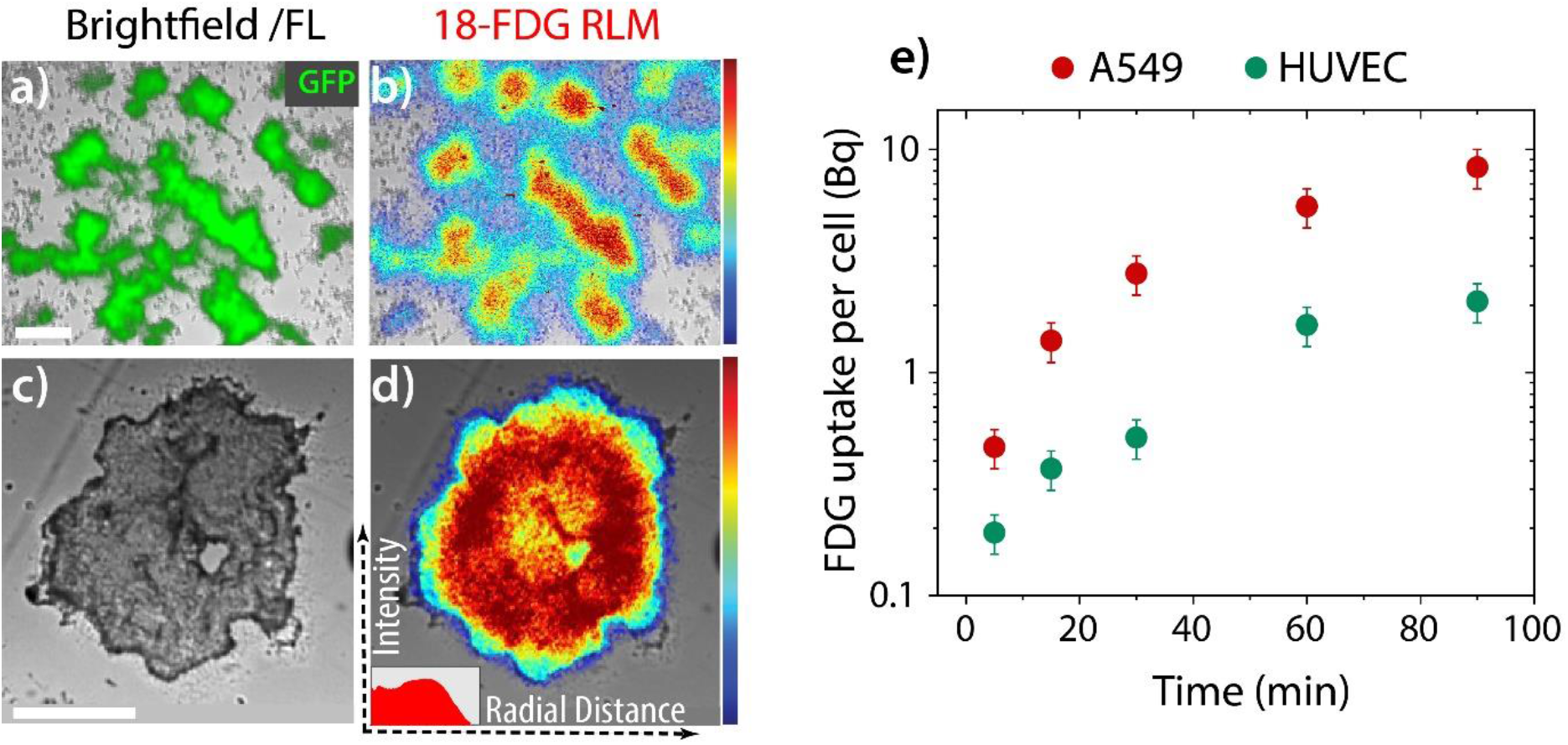
Characterization of FDG uptake in A549 and HUVEC cell cultures. (a) Brightfield and fluorescence overlay and (b) RLM imaging of HUVEC 3D culture in GelMA. The FDG and GFP images show a high degree of correlation with each other. (c-d) Brightfield and RLM images show high uptake of FDG in A549 spheroid. Inset: The heterogeneous FDG accumulation along the radial axis. Scale bar: 400 μm. (e) FDG uptake kinetics of A549 and HUVEC cell lines, measured by gamma counting. Error bars represent the uncertainty (standard deviation) during cell counting.

Additionally, the low FDG background in the RLM image confirms that unbound FDG is efficiently washed off from the hydrogel.

We then imaged FDG uptake in tumor spheroids. Due to their greater avidity for glucose, the spheroids could be imaged by the EMCCD using exposure as short as 3-5 s (Figure 3c, d). The radial distribution of accumulated FDG along the spheroid radius shows that glucose diffusion and metabolism are heterogeneous. This inhomogeneity might be correlated with the hypoxic area of the spheroid and limited penetration of FDG into the spheroid, but it was not confirmed in our study since we did not image hypoxia and FDG uptake in the same specimens. Finally, to characterize the intrinsic glycolytic rate, we measured FDG uptake kinetics by HUVEC and A549 cells cultured on a standard cell-culture dish. Comparing the two cell lines, we found that while they are both highly glycolytic, A549 cells accumulated over two-fold more FDG than HUVECs on a per-cell basis (Figure 3e).

Finally, we performed FDG imaging of co-cultured A549 spheroids (tumor) and HUVEC microvascular networks (stroma). The tumor spheroid shows a high concentration of FDG (Figure 4a-c) as expected from previous measurements. However, a significant amount of FDG was also retained in the stromal compartment of the co-culture. In addition to the uptake from HUVEC cells, the extracellular matrix too contributed to accumulating a small fraction of FDG in the tumor microenvironment. This model is relevant in many solid tumors where a large ECM-rich stromal component plays a key role in tumor maintenance. The co-culture with the smaller spheroids (Figure 4d-4f) was a more representative model of such tumors. The differential uptake of FDG (Figure 4g) shows that the tumor stroma in our model is metabolically active and largely FDG avid and could significantly contribute to the overall radioactivity or PET signal in the patient tumor.

**Figure 4:**
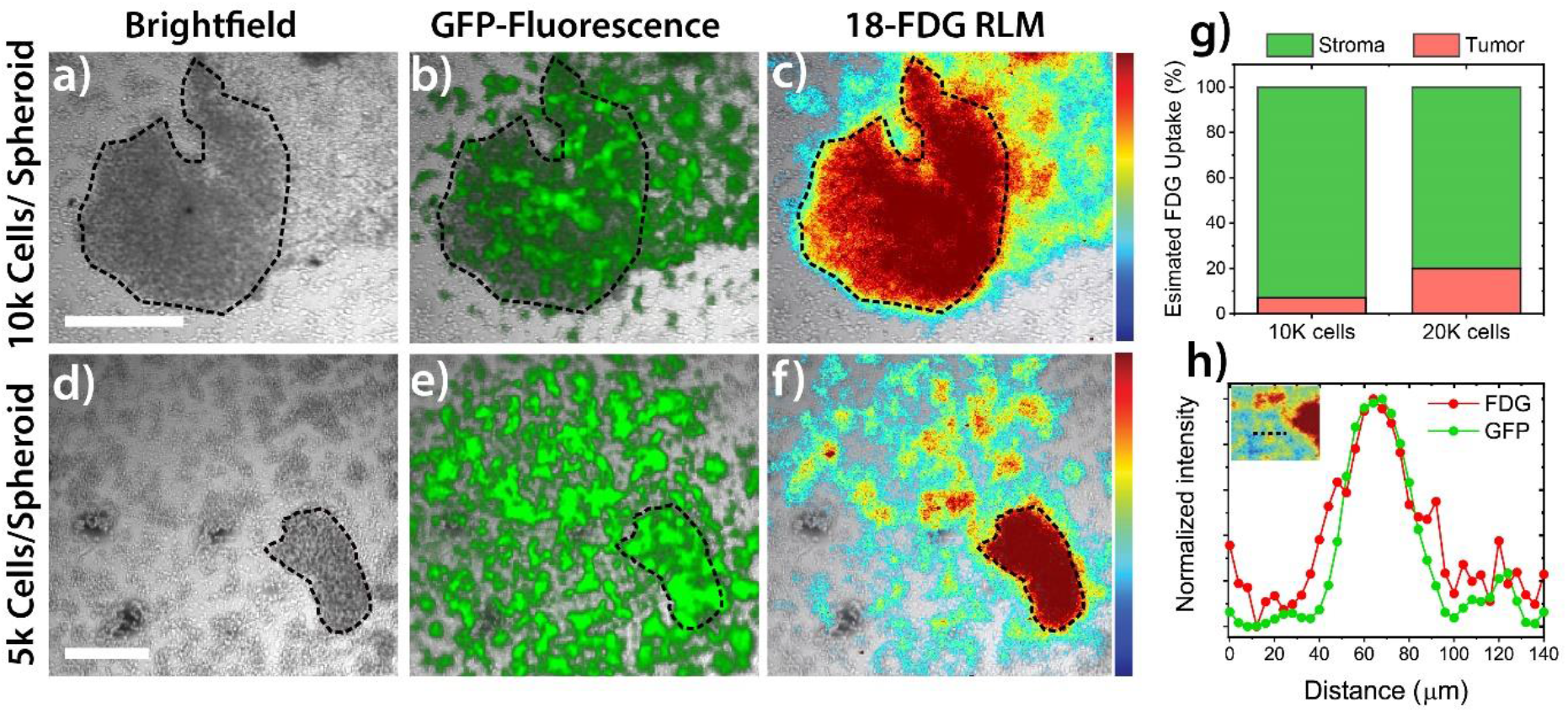
Metabolic imaging of tumor-stroma co-culture. (a, d) Brightfield, (b, e) GFP-fluorescence overlay and (b, f) RLM co-imaging of HUVEC-A549 co-culture culture in GelMA with two different spheroid sizes. RLM imaging shows the relative distribution of FDG. The accumulation of FDG in HUVEC can be visualized by spatially correlating with the GFP fluorescence (green). Scale bar: 0.5 mm (g) A comparative analysis of the % RLM intensity from the tumor and stromal compartment with two different spheroid size. (h) Co-registration of FDG and GFP signal intensity along a line-profile across an endothelial sprout.

### The limit of resolution

We previously demonstrated spatial resolution 20-30 μm using RLM for 2D cell culture imaging [9]. In this study, RLM is applied to thicker 3D tissue-engineered models, which may reduce the achievable spatial resolution due to the increase in positron travel. We found that RLM images show excellent spatial resolution and demonstrate the ability to resolve details much finer than those seen in a PET scan. For instance, the intricate HUVEC network was clearly visible in our images. To estimate the approximate limit of resolution in this study, we plotted a line profile across a ~100 μm wide endothelial sprout (Figure 4h). The line profile of the FDG signal overlapped visibly with the GFP fluorescence signal. The full-width half-maxima of the Gaussian profile was found to be 32±3 μm. This range is close to the dimension of individual cells, indicating >100 times higher resolving power than clinical or preclinical PET imaging techniques.

## Discussion

In this study, we have developed a basic 3D tumor-stroma model using human lung adenocarcinoma and endothelial cells. We demonstrated that, in such a model, RLM can be effectively used for high-resolution imaging of glucose metabolism within the 3D tumor and stromal compartments. The high uptake of glucose by endothelial cells contributed significantly towards the overall FDG signal from the tumor model. This new technique can be used to image most *in vitro* cancer models, including surgical tissue slices, organoids, and tumor biopsies, using relevant clinical PET radioisotopes.

This study demonstrates that RLM imaging can be performed on the live 3D-cultured specimen, in spite of their increased thickness. This is a crucial capability since these thicker specimens are more representative of tumor physiology, but one challenge is that spatial resolution and sensitivity are degraded due to the propagation and scattering of the positrons. For maximal sensitivity, RLM does not collimate the incoming positrons, thus spatial information is derived from the proximity of the source to the scintillator. In a previous computational study [14], we found that the spatial resolution of RLM degrades at a rate of ~3 μm FWHM per 1 μm distance as the source is moved away from the scintillator. Additionally, the signal intensity from a point source seen by the scintillator decreases according to the inverse square law (beam divergence) and positron range. Accordingly, the resulting effect is that image resolution and sensitivity is depth-dependent. These effects could potentially be corrected if the depth of the sources was known.

It has been postulated that the angiogenic switch, which is characterized by rapid blood vessel sprouting and tumor vascularization, requires a metabolic switch as well. The ECs are known to be highly glycolytic and meet up to 85% of their ATP demand only through the glycolysis pathway [5, 15]. It is not clear why endothelial cells prefer a less energy-efficient pathway of ATP production. Multiple hypotheses exist, including the need of transferring more oxygen to perivascular tissue, or an adaptation to the hypoxic avascular tissue environment during vascularization, and thus preferring oxygen-independent metabolic route. Moreover, it has been shown that glycolysis intermediates are utilized by ECs for anabolic biosynthesis of macromolecules, angiogenesis signaling, migration, proliferation, and survival. These hypotheses can be tested by RLM in future studies.

Similarly, cancer-associated fibroblasts which closely nurture and promote the growth of cancer cells are thought to develop metabolic pathways complementary to those of cancer cells, setting up a tumor-stroma metabolic loop. Studies have suggested that cancer-associated fibroblasts are often glycolytic, especially for invasive cancers, and therefore may contribute significantly to the total tumor uptake of FDG during PET imaging [3, 16]. These fibroblasts also promote angiogenesis by secreting growth factors, recruiting and proliferating endothelial cells. These metabolic adaptations of the stroma can be efficiently investigated at high resolution in appropriate *in vitro* models utilizing the RLM imaging technique.

A growing number of clinical PET tracers have been developed to detect specific alterations of the tumor microenvironment. For instance, fibroblasts activation proteins are displayed by tumor-associated fibroblasts in a variety of cancers and can be imaged using PET. In addition, the prostate-specific membrane antigen is commonly expressed on the neovasculature of some tumors. Tumor-infiltrating lymphocytes and activated T-cells can be also be imaged using various PET tracers. In this respect, engineered tumor microenvironments that contain stromal and/or immune cells provide an attractive platform to develop and investigate PET tracers. RLM can image commonly used PET radioisotopes including ^18^F, ^68^Ga, ^64^Cu, ^124^I, and ^11^C. As only a small section of the scintillator is in focus during imaging, the larger positron range of some of the above isotopes does not degrade the spatial resolution in RLM. Moreover, therapeutic alpha and beta radiation could be imaged for microdosimetry evaluation of radiopharmaceutical therapy.

With the advancement of tissue engineering and organoid technology, appropriate tissue-construct models can be tailored to evaluate and screen specific PET tracers with high-resolution using our imaging technique. For instance, using FDG, we have recently shown that patient-derived head & neck tumor organoids recapitulate the metabolic heterogeneity of the original tumor [17]. We also tested the feasibility of screening for patient-specific drug resistance, and aim to extend this technique for high-throughput screening. In addition, other in vitro models are being developed to better mimic human physiology in the lab [18, 19]. Perfused organ-on-chips can be potentially coupled with RLM for high-resolution dynamic imaging and of PET tracer *in vitro*. For instance, a brain-on-chip could be used to test new brain imaging tracers, whereas kidney-on-chip and liver-on-chip could be utilized to evaluate excretion and metabolism of PET tracers. Similarly, new targeted PET tracers can be evaluated in cohorts of patient-specific cancer-chips constructed with patient-derived tumor tissue.

In conclusion, RLM creates new opportunities for using PET radiopharmaceuticals to image advanced *in vitro* cancer models and other tissue-engineered constructs. As a first demonstration, we used FDG uptake to estimate the metabolic flux of glucose in the endothelial tumor stroma in a 3D co-culture model. With recent advances on *in vitro* modeling of the tumor microenvironment, RLM provides a new approach to investigate the uptake and distribution of various PET tracer using *in vitro* models and achieve a new level of understanding of these PET tracers, beyond the limitations imposed by the coarse spatial resolution of PET imaging.

## References

1. Vander Heiden MG, Cantley LC, Thompson CB. Understanding the Warburg Effect: The Metabolic Requirements of Cell Proliferation. Science. 2009;324:1029. doi:10.1126/science.1160809.

2. Moses WW. Fundamental limits of spatial resolution in PET. Nuclear Instruments and Methods in Physics Research Section A: Accelerators, Spectrometers, Detectors and Associated Equipment. 2011;648:S236–S40. doi:https://doi.org/10.1016/j.nima.2010.11.092.

3. Avagliano A, Granato G, Ruocco MR, Romano V, Belviso I, Carfora A, et al. Metabolic Reprogramming of Cancer Associated Fibroblasts: The Slavery of Stromal Fibroblasts. BioMed Research International. 2018;2018:6075403. doi:10.1155/2018/6075403.

4. Fitzgerald G, Soro-Arnaiz I, De Bock K. The Warburg Effect in Endothelial Cells and its Potential as an Anti-angiogenic Target in Cancer. Frontiers in Cell and Developmental Biology. 2018;6. doi:10.3389/fcell.2018.00100.

5. Ghesquière B, Wong BW, Kuchnio A, Carmeliet P. Metabolism of stromal and immune cells in health and disease. Nature. 2014;511:167–76.

6. Gómez V, Eykyn TR, Mustapha R, Flores-Borja F, Male V, Barber PR, et al. Breast cancer-associated macrophages promote tumorigenesis by suppressing succinate dehydrogenase in tumor cells. Science signaling. 2020;13.

7. Kim TJ, Türkcan S, Pratx G. Modular low-light microscope for imaging cellular bioluminescence and radioluminescence. Nature protocols. 2017;12:1055.

8. Liu Z, Lan X. Microfluidic radiobioassays: a radiometric detection tool for understanding cellular physiology and pharmacokinetics. Lab on a Chip. 2019;19:2315–39. doi:10.1039/C9LC00159J.

9. Pratx G, Chen K, Conroy Sun LM, Carpenter CM, Olcott PD, Xing L. Radioluminescence microscopy: measuring the heterogeneous uptake of radiotracers in single living cells. PloS one. 2012;7.

10. Sengupta D, Pratx G. Radioluminescence Microscopy: A Quantitative Method for Radioisotopic Imaging of Metabolic Fluxes in Living Cancer Cells. In: Haznadar M, editor. Cancer Metabolism: Methods and Protocols. New York, NY: Springer New York; 2019. p. 45–53.

11. Nichol JW, Koshy ST, Bae H, Hwang CM, Yamanlar S, Khademhosseini A. Cell-laden microengineered gelatin methacrylate hydrogels. Biomaterials. 2010;31:5536–44.

12. Pratx G, Chen K, Sun C, Axente M, Sasportas L, Carpenter C, et al. High-Resolution Radioluminescence Microscopy of 18F-FDG Uptake by Reconstructing the β-Ionization Track. Journal of Nuclear Medicine. 2013;54:1841–6. doi:10.2967/jnumed.112.113365.

13. Anada T, Pan C-C, Stahl AM, Mori S, Fukuda J, Suzuki O, et al. Vascularized bone-mimetic hydrogel constructs by 3D bioprinting to promote osteogenesis and angiogenesis. International journal of molecular sciences. 2019;20:1096.

14. Wang H, Ma Y, Pratx G, Xing L. Toward real-time Monte Carlo simulation using a commercial cloud computing infrastructure. Physics in Medicine and Biology. 2011;56:N175–N81. doi:10.1088/0031-9155/56/17/n02.

15. De Bock K, Georgiadou M, Schoors S, Kuchnio A, Wong Brian W, Cantelmo Anna R, et al. Role of PFKFB3-Driven Glycolysis in Vessel Sprouting. Cell. 2013;154:651–63. doi:https://doi.org/10.1016/j.cell.2013.06.037.

16. Shangguan C, Gan G, Zhang J, Wu J, Miao Y, Zhang M, et al. Cancer-associated fibroblasts enhance tumor 18F-FDG uptake and contribute to the intratumor heterogeneity of PET-CT. Theranostics. 2018;8:1376.

17. Khan S, Shin JH, Ferri V, Cheng N, Noel JE, Kuo C, et al. High-resolution positron emission microscopy of patient-derived tumor organoids. bioRxiv. 2020:2020.07.28.220343. doi:10.1101/2020.07.28.220343.

18. Tuveson D, Clevers H. Cancer modeling meets human organoid technology. Science. 2019;364:952–5.

19. Low LA, Mummery C, Berridge BR, Austin CP, Tagle DA. Organs-on-chips: into the next decade. Nature Reviews Drug Discovery. 2020. doi:10.1038/s41573-020-0079-3.

